# An updated vocal repertoire of wild adult bonobos (*Pan paniscus*)

**DOI:** 10.1101/2025.01.23.634282

**Authors:** Franziska Wegdell, Isaac Schamberg, Mélissa Berthet, Yannik Rothacher, Volker Dellwo, Martin Surbeck, Simon Townsend

**Author notes:** Corresponding author (FW). These authors contributed equally (Martin Surbeck, Simon Townsend).

## Abstract

While research over the last 20 years has shed important light on the vocal behaviour of our closest living relatives, bonobos and chimpanzees, a quantitative description of their full vocal repertoire is absent. Such data are critical for a holistic understanding of a species’ communication system and unpacking how these systems compare more broadly with other primate and non-primate species. Here we make key progress by providing the first quantitative *Pan* vocal repertoire, specifically for wild bonobos. Using data comprising over 1500 calls from 53 adult individuals collected over 33 months, we employ machine-learning-based random forest analyses and describe 11 acoustically distinguishable call types. We discuss issues associated with resolving vocal repertoires from wild data in great apes and highlight potential future approaches to further capture the complexity of the bonobo vocal system.

## Introduction

Over the last 20 years, considerable research attention has focused on the vocal behaviour of our closest-living relatives, non-human great apes, not least given the insights such findings can provide into the evolution of our own communication system: language. Unsurprisingly, a significant proportion of work to date has focused on our closest cousins, the *Pan* lineage (chimpanzees (*Pan troglodytes*) and bonobos (*Pan paniscus*)). Research in chimpanzees has demonstrated that they use calls and call combinations flexibly across contexts (Bortolato et al. 2023; Crockford 2019); possess a degree of plasticity within call production (Mitani and Gros-Louis 1998; Watson et al. 2015); deploy calls voluntarily (Crockford et al. 2012; Girard-Buttoz et al. 2020; Kalan and Boesch 2015; Schel et al. 2013; Townsend et al. 2017; Townsend, Deschner, and Zuberbühler 2008); produce call combinations (Crockford and Boesch 2005; Girard-Buttoz et al. 2022a; Leroux et al. 2021); extract meaning from such combinations (Leroux et al. 2023) and the call order can differ between populations (Girard-Buttoz et al. 2022b). Similar findings have been demonstrated in bonobos; individuals also deploy calls flexibly across contexts (Genty et al. 2014), deploy calls voluntarily (Girard-Buttoz et al. 2020) use call combinations (Schamberg et al. 2016; 2017) and can derive meaning from them (Clay and Zuberbühler 2009, 2011). In addition, certain calls demonstrate functional flexibility (i.e. where calls can be used across contexts characterized by different emotional valence) (Clay, Archbold, and Zuberbühler 2015) and populations differ in call usage (Schamberg et al. 2023; 2024). Despite considerable progress in decomposing the complexity of *Pan* vocal systems, in both wild chimpanzees and bonobos, only qualitative descriptions of their vocal repertoires exist. Potential reasons for this include, but are not limited to, the fact that *Pan* vocal repertoires are graded (Crockford 2019), where calls overlap in a continuous acoustic space and quantitative approaches to resolving a call repertoire necessarily require a large and robust sample size. Here, using data collected over 33 months, we make key progress on this issue by providing the first quantitative *Pan* vocal repertoire using calls specifically from wild bonobos.

Previous attempts to describe the bonobo vocal repertoire do exist. First, de Waal (1988) took a qualitative approach (with calls being classed into call type categories by human ear or by visually assessing spectrograms) recording ten captive, mainly immature individuals and described 17 call types. Hopkins & Savage-Rumbaugh (1991) described 10 call types for group-living captive bonobos and 14 for a human-reared bonobo. Almost a decade later, Bermejo & Omedes (1999) followed this up and published the only vocal repertoire on wild bonobos to date, which again qualitatively describes 15 call types. More recently, Keenan and colleagues (2020) and Arnaud and colleagues (2023) have revisited the vocal repertoire of captive bonobos and, although quantitative methods were employed, only a subset of calls from the repertoire (namely *high hoots, barks, soft barks, peep-yelps* and *peeps*) were considered.

Whilst qualitative repertoires serve as an important entry point facilitating the investigation into non-human animal vocal behaviour, they are unavoidably subjective and susceptible to bias since they rely purely on human visual and auditory discrimination. One consequence of this subjective approach is that various researchers seem to carve up the repertoire of the same species differently. Bonobos are no exception and hence much inconsistency between studies regarding the naming and definition of call types exits. Bermejo and Omedes (1999) for example used the term *barks* for calls that other studies (Hohmann and Fruth 1994; Schamberg et al. 2017; De Waal 1988) termed *high hoots*; and did not distinguish *contest hoots* (described in de Waal (1988) and Genty et al. (2014)) from other types of hoots. As another example, *soft barks* were described by Bermejo and Omedes (1999) and Keenan (2020), but Clay and Zuberbühler (2009) used the label *food barks*, and de Waal (1980) did not describe this call type at all. In order to avoid issues with subjective categorisation, quantitative vocal repertoires, whereby calls are classified based on their acoustic features, represent a constructive move forward. Such approaches further allow specific questions that were previously intractable to be addressed. For example, quantitative repertoires can be compared more reliably across individuals, groups, populations and species. Additionally, quantitative repertoires can facilitate and optimize conservation-driven questions such as passive acoustic monitoring-based population surveys where knowledge regarding the vocal repertoire of a species is needed (Sills and Reichmuth 2022) and which are more efficient than camera traps in detecting primates (Crunchant et al. 2020).

Quantitative approaches to describe vocal repertoires range from traditional acoustic analysis and discrimination-based statistical analyses (Hending, Seiler, and Stanger-Hall 2020; Maretti et al. 2010; Salmi, Hammerschmidt, and Doran-Sheehy 2013), to more state-of-the-art machine learning-based methods where supervised or unsupervised algorithms cluster calls into defined or undefined categories, respectively (Keen et al. 2021; Keenan et al. 2020; Sainburg, Thielk, and Gentner 2020). Here we attempt to quantitatively assess and validate the hitherto described call types of the vocal repertoire of wild adult bonobos specifically using machine learning-based supervised random forest classifying algorithms.

## Material and methods

### Ethics statement

Ethical permission to conduct this non-invasive study was granted by the Institut Congolais pour la Conservations de la Nature and the Ministry of Research and Technology of the Democratic Republic of the Congo. This study is in line with the ethical guidelines of the former Department of Primatology at the Max-Planck-Institute for Evolutionary Anthropology and the guidelines of the American Society of Primatologists for the ethical treatment of non-human primates. We also adhered to the best practice guidelines for health monitoring and disease control in great ape populations (Gilardi et al. 2015): researchers underwent quarantine, wore masks during data collection and maintained a 7m distance to the bonobos. Access to the Kokolopori Bonobo Reserve was granted by the villages of Bolamba, Yete, Yomboli, and Yasalakose. Additional information regarding the ethical, cultural, and scientific considerations specific to inclusivity in global research is included in the Supporting Information (S7 Questionnaire)

### Study site and subjects

We recorded vocalisations using *ad libitum* and focal sampling methods (Altmann 1974) for 33 months over a period of 10 years (2011-2022) from habituated wild adult bonobos. Subjects were recorded at two field sites in the Democratic Republic of Congo, the Kokolopori Bonobo Reserve (Surbeck, Coxe, and Lokasola 2017) and the Luikotale field site (Hohmann and Fruth 2003) from a total of four bonobo communities. At the Kokolopori Bonobo Reserve, FW, MB and IS collected data over 20 months; at Luikotale, IS collected data over a period of 13 months. We recorded a total of 1509 vocalisations with broadly similar numbers of calls from female and male individuals (804 and 705 respectively). Individuals from one community in Luikotale were recorded and contributed 672 calls and in Kokolopori individuals from the Kokoalongo, Ekalakala and Fekako community contributed 526, 241 and 70 calls, respectively. On average we used 28 (range 1-145) calls per individual from 53 adult individuals for further analyses.

### Data collection

We recorded all vocalisations with a 44.1kHz sampling frequency and a 16-bit amplitude resolution with Marantz PMD 660 digital recorders and Sennheiser directional microphones (K6 power module, ME66 recording head and Rycote-Softie windscreen) at a distance of 7-10m. For each vocalisation we noted the date and ID of the caller.

### Data preparation

Only calls for which the caller was known were included in the analysis. We visually inspected spectrograms using Adobe Audition 2020 software (v. 13.0.13.46, Adobe Systems Inc., San Jose, CA, U.S.A.) with a Hamming window and a 256-frequency step. We extracted non-overlapped call units of sufficient quality (e.g. minimal background noise, no clipping) for further analysis. We categorized the calls into 15 “original call types” based on spectrographic and acoustic descriptions from previously published repertoires (Bermejo and Omedes 1999; Keenan et al. 2020; De Waal 1988). For an overview of the sample size for each of the 15 original call types per individual, see S1 Table.

### Data analysis

#### Acoustic measurements

Using the R (v. 4.3.1 (R Core Team 2024)) package warbleR (Araya-Salas and Smith-Vidaurre 2017) we used a 200 Hz high-pass filter and a 4000 Hz low-pass filter to remove low and high frequency noise (e.g. cicadas). For each call, we automatically extracted 26 time- and frequency-related acoustic parameters commonly used in bioacoustic analyses (see S2 Table for an exact description of all parameters) using the package warbleR (Araya-Salas and Smith-Vidaurre 2017) (see supplementary R code for more information). Following Keen et al. 2021, we also calculated a pairwise distance matrix using dynamic time warping. We used classical multi-dimensional scaling (MDS) to translate the matrix into a five-dimensional space, and used the axis coordinates for each sample as additional call metrics (i.e., five dynamic time wrapping MDS coordinates per call). This resulted in a total of 31 automatically extracted parameters for each call. To mitigate errors from automatic extraction of acoustic parameters whilst simultaneously retaining the calls in the dataset, we identified outliers by calculating a z-score for each acoustic parameter. Calls for which the absolute z-score was above 3.29 for a specific acoustic parameter were considered outliers: these values were replaced by the median of that acoustic parameter, following Santhanam and Padmavathi (2014). In total we identified 305 parameter values considered to be outliers from 148 calls.

#### Call type classification

An unsupervised classification method (i.e., which detects underlying structure within unlabelled data) would be an objective approach and ensure a robust repertoire, but relies on substantially larger datasets. In a pilot step, we conducted an unsupervised random forest analysis, which could only categorise calls into two call types (see S3 Analysis). Previous work on bonobo vocal behaviour (e.g. Arnaud et al. 2023; Bermejo and Omedes 1999; Clay, Archbold, and Zuberbühler 2015; Clay and Zuberbühler 2009, 2011; Genty et al. 2014; Hohmann and Fruth 1994; Keenan et al. 2020; Schamberg et al. 2017, 2023; De Waal 1988), indicates bonobo vocalizations fall into more than two categories. As such, this result probably does not reflect acoustic differences between the calls, but is rather a consequence of the limitations of the methodology - namely a relatively small dataset for the amount of call types and the inherent gradedness of the bonobo vocal system. Since a large enough dataset necessary to implement unsupervised analyses is not available, supervised approaches are, to our knowledge, the only relevant alternative.

We followed the methods laid out in Keen et al. (2021) for using supervised random forest analyses to assess the robustness of call classification. The random forest analysis is a machine learning method which creates a set of decision trees (Kotsiantis 2013) wherein, at each node of each tree, the data is divided into two classes using a random subset of the acoustic parameters (Breiman 2001; Rothacher and Strobl 2023; Strobl, Malley, and Tutz 2009). Each datapoint is then assigned a call type based on the category chosen by the majority of trees. Finally, the data is classified as well as possible into the given categories (the “original call types”). We used the randomForest R package (Liaw and Wiener 2002) to implement the supervised random forest with 1000 decision trees and five (i.e., the square root of the total number of features) randomly selected acoustic parameters at each split (for more details, see Breiman 2001 and Liaw and Wiener 2002).

Random forest classifiers are considered one of the best available classification methods, as they work better than other machine learning algorithms on small datasets and can better detect small differences between classes (see Carugati et al. 2024, Wieruka et al., Biorxiv preprint). For this reason this approach has been commonly and successfully used in similar studies to describe vocal repertoires (e.g. Keen at al. (2021).

We deemed a call type to be reliably classified, and hence acoustically discriminable, if the random forest was able to correctly classify a plurality of the calls within a given call type as the “original call type” (i.e., the count of calls labelled as the initial call type by the random forest was the highest). If the random forest incorrectly classified the plurality of calls within a given call type, we did not consider it acoustically distinct and the initial putative call type was relabelled as the call type that the random forest most often classified it as. To help clarify our approach, consider a hypothetical dataset with three initial putative call types – A, B, and C – each of which has 100 datapoints. After training a random forest model on the three putative call types, the model correctly classified 20/100 calls labelled call type A, and incorrectly classified the other 80/100 (50/100 were classified as call type B and 30/100 were classified as call type C). The model correctly classified 100% of calls labelled call type B and call type C. According to our criterion, call types B and C would be considered valid call types, but call type A would not be considered acoustically discriminable because the model classified it as a different call type in a plurality of cases. After this result, all calls labelled call type A would be relabelled call type B because it was classified as call type B in the plurality of cases.

Finally, to estimate the significance of the overall classification we used a two-tailed binomial test with a level of chance corresponding to the number of call types to be classified (i.e. 1/15=0.067).

#### Data visualization

We employed a non-linear dimensionality reduction algorithm (i.e. t-Distributed Stochastic Neighbor Embedding (t-SNE)) to visualize the clustering of the call types of the updated vocal repertoire. Specifically, a t-SNE provides a visual representation of how acoustically similar calls cluster together in a 2-D space (van der Maaten 2014; van der Maaten and Hinton 2008).

## Results

The random forest agreed with the “original call types” on the classification of 868 of the 1509 calls (57.52% of the calls), a rate significantly exceeding the probability of a call being assigned to one of the 15 call types by chance (1/15*100=6.67%, binomial test p=<0.001). Accordingly, the probability of being incorrectly classified (the out-of-bag error rate) was 42.48%. The classification error varied with the call types (Table 1), with some call types being more accurately classified (e.g. *high hoots, screams* and *laughter*) and others less so (e.g. *barks, scream bark*). Eleven of the 15 call types met our criterion to be considered a reliably acoustically discriminable vocalisation (Table 2; for more information on the influence each acoustic variable had on the random forest see S4 Fig). Follow-up t-SNE-based visualisation of the data indicates that whilst these 11 acoustically discriminable call types could be reliably identified, the bonobo vocal repertoire is still highly graded (Fig 1). We provide an overview of the 11 defined call types with an accompanying spectrogram and description of how to identify them (Fig 2) and the mean and range for common parameters (Table 3).

**Table 1.** Confusion matrix illustrating the classification of call types by the random forest clustering analysis.

| Call type | Predicted group membership |  |  |  |  |  |  |  |  |  |  |  |  |  |  | Classification error |
| --- | --- | --- | --- | --- | --- | --- | --- | --- | --- | --- | --- | --- | --- | --- | --- | --- |
|  | *high hoot* | *scream* | *grunt* | *peep* | *laughter* | *low hoot* | *whistle* | *contest-hoot* | *pant grunt* | *yelp* | *peep yelp* | *scream bark* | *wieew bark* | *bark* | *soft bark* |  |
| *high hoot* | **242** | 3 | 0 | 0 | 0 | 0 | 3 | 2 | 0 | 4 | 12 | 0 | 7 | 10 | 2 | 0.151 |
| *scream* | 11 | **56** | 0 | 3 | 0 | 0 | 8 | 1 | 0 | 1 | 0 | 3 | 0 | 3 | 0 | 0.349 |
| *grunt* | 0 | 0 | **99** | 6 | 2 | 8 | 0 | 0 | 8 | 4 | 8 | 0 | 0 | 0 | 0 | 0.267 |
| *peep* | 5 | 0 | 9 | **105** | 0 | 0 | 3 | 1 | 2 | 6 | 18 | 0 | 0 | 0 | 2 | 0.305 |
| *laughter* | 1 | 0 | 6 | 0 | **19** | 0 | 1 | 0 | 0 | 1 | 0 | 0 | 1 | 0 | 0 | 0.345 |
| *low hoot* | 2 | 0 | 21 | 0 | 0 | **50** | 0 | 0 | 0 | 1 | 0 | 0 | 2 | 0 | 0 | 0.342 |
| *whistle* | 12 | 6 | 0 | 4 | 0 | 0 | **58** | 2 | 0 | 2 | 4 | 0 | 0 | 2 | 1 | 0.363 |
| *contest hoot* | 7 | 0 | 0 | 3 | 0 | 0 | 1 | **36** | 0 | 1 | 4 | 2 | 0 | 1 | 3 | 0.379 |
| *pant grunt* | 0 | 0 | 6 | 1 | 0 | 3 | 0 | 0 | **32** | 1 | 9 | 0 | 0 | 0 | 1 | 0.396 |
| *yelp* | 8 | 0 | 13 | 13 | 0 | 3 | 3 | 3 | 1 | **44** | 18 | 0 | 0 | 0 | 1 | 0.589 |
| *peep yelp* | 14 | 0 | 15 | 26 | 0 | 2 | 3 | 1 | 2 | 14 | **72** | 0 | 0 | 3 | 5 | 0.541 |
| *scream bark* | 13 | 4 | 0 | 0 | 0 | 0 | 4 | 5 | 0 | 0 | 2 | **11** | 0 | 1 | 0 | 0.725 |
| *wieew bark* | 32 | 1 | 2 | 0 | 0 | 1 | 0 | 1 | 0 | 2 | 3 | 0 | **15** | 0 | 1 | 0.741 |
| *bark* | 41 | 2 | 3 | 2 | 0 | 1 | 6 | 1 | 1 | 0 | 16 | 1 | 1 | **25** | 4 | 0.760 |
| *soft bark* | 22 | 0 | 5 | 3 | 0 | 1 | 4 | 10 | 3 | 2 | 16 | 1 | 1 | 7 | **4** | 0.950 |

*The random forest analysis classified call types based on their acoustic features. Bold numbers in the diagonal show the* *absolute number of calls being classified correctly as an original call type. The non-bold cells show the absolute number of* *calls that were categorized as another call type. We deemed a call type to be reliably discriminated when it was consistently* *classified by the random forest (i.e., the count of calls labelled as the original call type by the random forest was the highest).* *The sample size of the “original call types” is the sum of the numbers in one row. Calls below the black line are those that are* *classified more as other call types than themselves and hence merged with the call type they were confused with most (see S.*

**Table 2.** Originally used call types and call types they were subsequently merged into, based on the random forest analysis.

| **Original Call type** | **Updated Vocal Repertoire** |
| --- | --- |
| *high hoot* | *high hoot* |
| *scream bark* |  |
| *wieew bark* |  |
| *bark* |  |
| *soft bark* |  |
| *scream* | *scream* |
| *grunt* | *grunt* |
| *peep* | *peep* |
| *laughter* | *laughter* |
| *low hoot* | *low hoot* |
| *whistle* | *whistle* |
| *contest hoot* | *contest hoot* |
| *pant grunt* | *pant grunt* |
| *yelp* | *yelp* |
| *peep yelp* | *peep yelp* |

**Table 3.** Mean and range of the parameters duration, peak frequency, dominant frequency and mean frequency for each call type.

| Call type | Duration (s) | Peak frequency (kHz) | Dominant frequency (kHz) | Mean frequency (kHz) |
| --- | --- | --- | --- | --- |
| *contest hoot* | $0.24 \pm 0.32$ | $2.32 \pm 1.98$ | $2.24 \pm 1.46$ | $2.31 \pm 1.2$ |
| *grunt* | $0.1 \pm 0.28$ | $0.75 \pm 2.85$ | $0.9 \pm 2.64$ | $1.59 \pm 1.78$ |
| *high hoot* | $0.25 \pm 0.63$ | $2.3 \pm 4.06$ | $2.03 \pm 2.59$ | $2.25 \pm 2.51$ |
| *laughter* | $0.14 \pm 0.36$ | $0.87 \pm 4.06$ | $1.12 \pm 2.45$ | $1.76 \pm 0.88$ |
| *low hoot* | $0.12 \pm 0.43$ | $0.55 \pm 1.12$ | $0.63 \pm 0.89$ | $1.2 \pm 1.31$ |
| *peep* | $0.14 \pm 0.47$ | $1.85 \pm 3.68$ | $1.97 \pm 3.19$ | $2.03 \pm 1.77$ |
| *pant grunt* | $0.08 \pm 0.12$ | $0.86 \pm 1.98$ | $0.99 \pm 0.88$ | $1.44 \pm 0.76$ |
| *peep-yelp* | $0.14 \pm 0.23$ | $1.55 \pm 4.06$ | $1.55 \pm 2.2$ | $1.96 \pm 1.72$ |
| *scream* | $0.58 \pm 1.24$ | $3.54 \pm 3.37$ | $3.37 \pm 2.65$ | $2.85 \pm 1.98$ |
| *whistle* | $0.5 \pm 1.47$ | $2.18 \pm 4.06$ | $2.27 \pm 1.9$ | $2.36 \pm 1.86$ |
| *yelp* | $0.15 \pm 0.48$ | $1.78 \pm 3.97$ | $1.85 \pm 1.5$ | $2.04 \pm 1.83$ |

**Fig 1.**
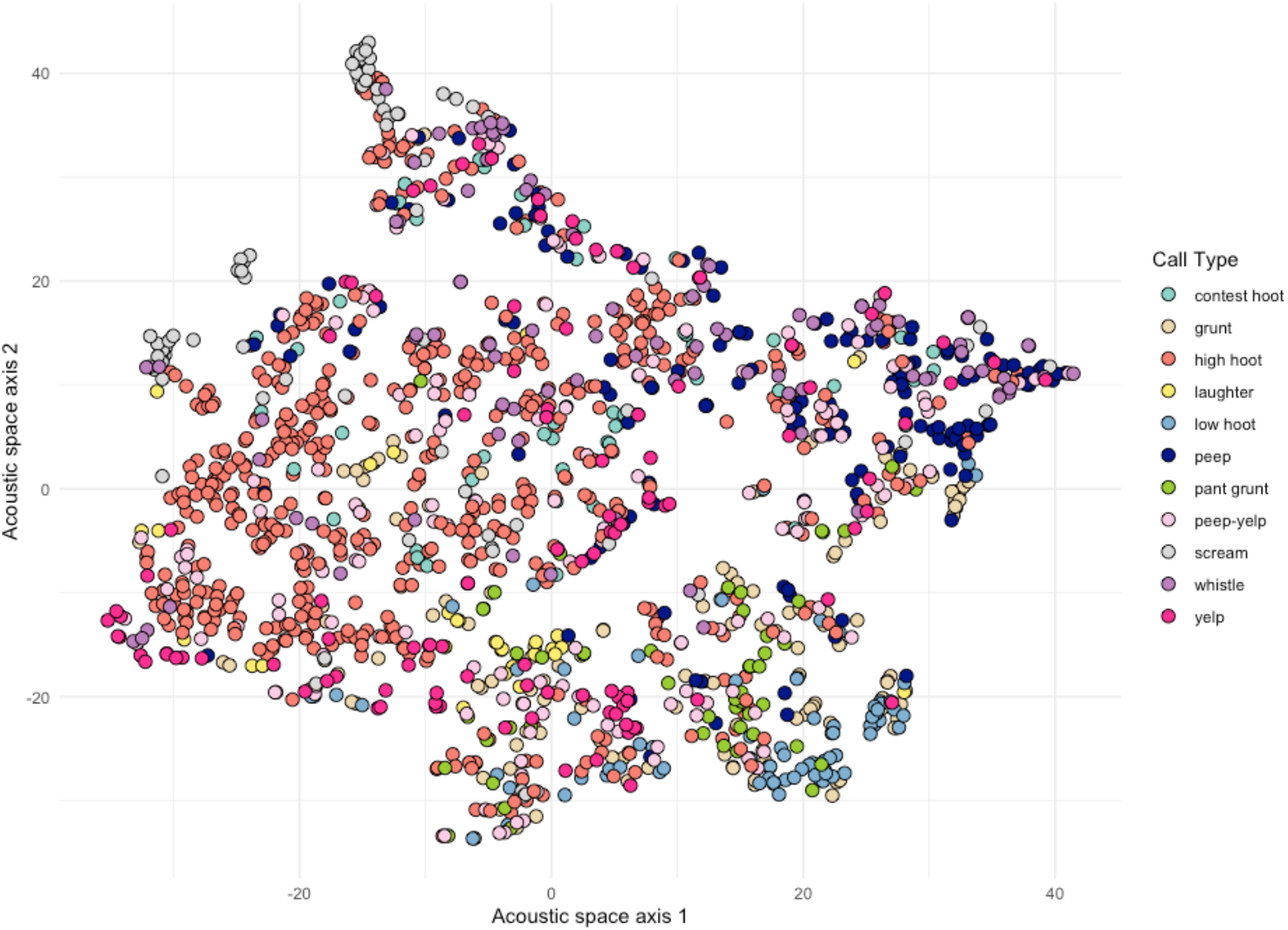
T-SNE scatterplot illustrating how the 11 call types from the updated vocal repertoire cluster together based on similarity in their acoustic structure. Each point in the scatterplot represents a call and the different colours of the points depict the different call types.

**Fig 2.**
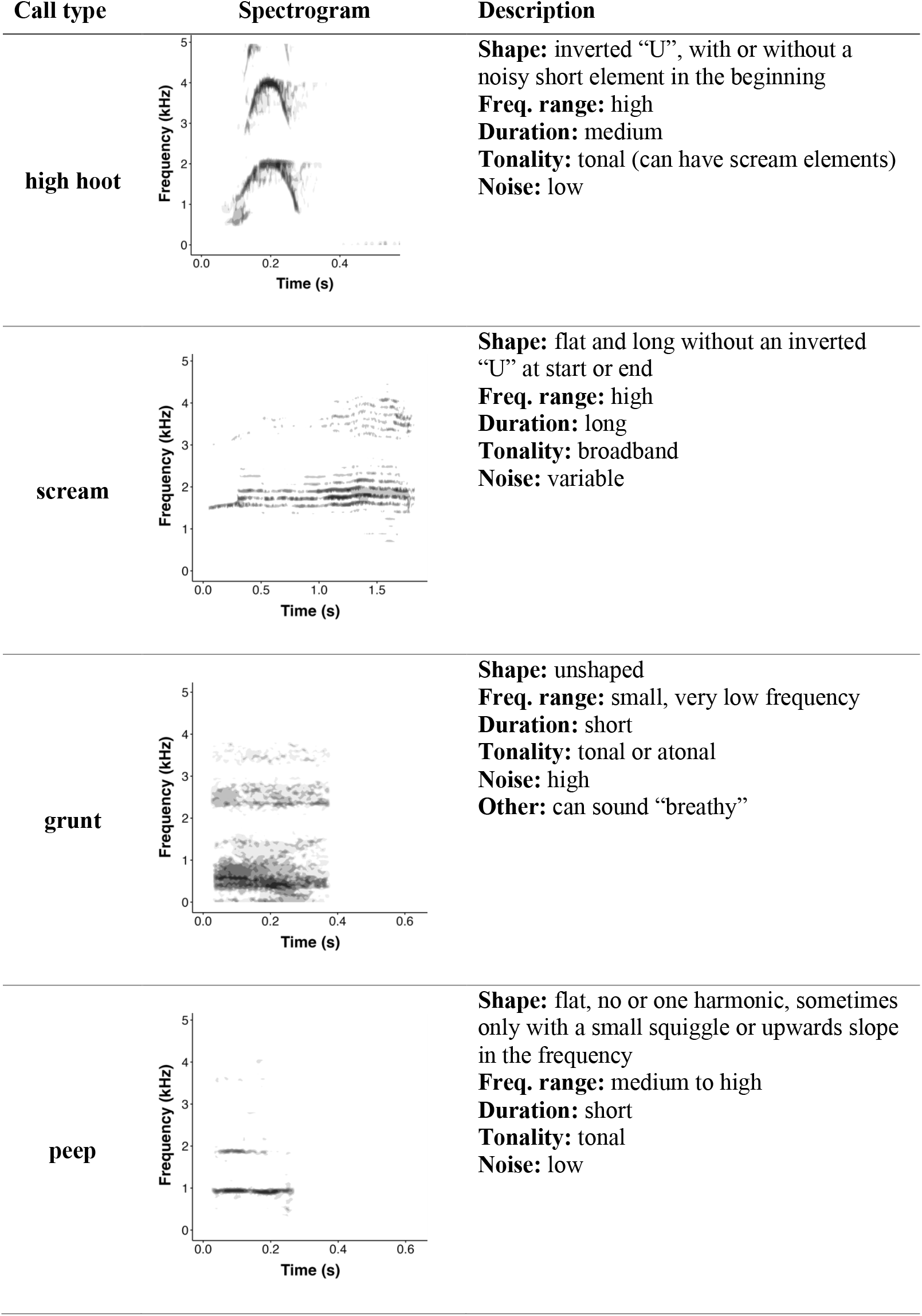
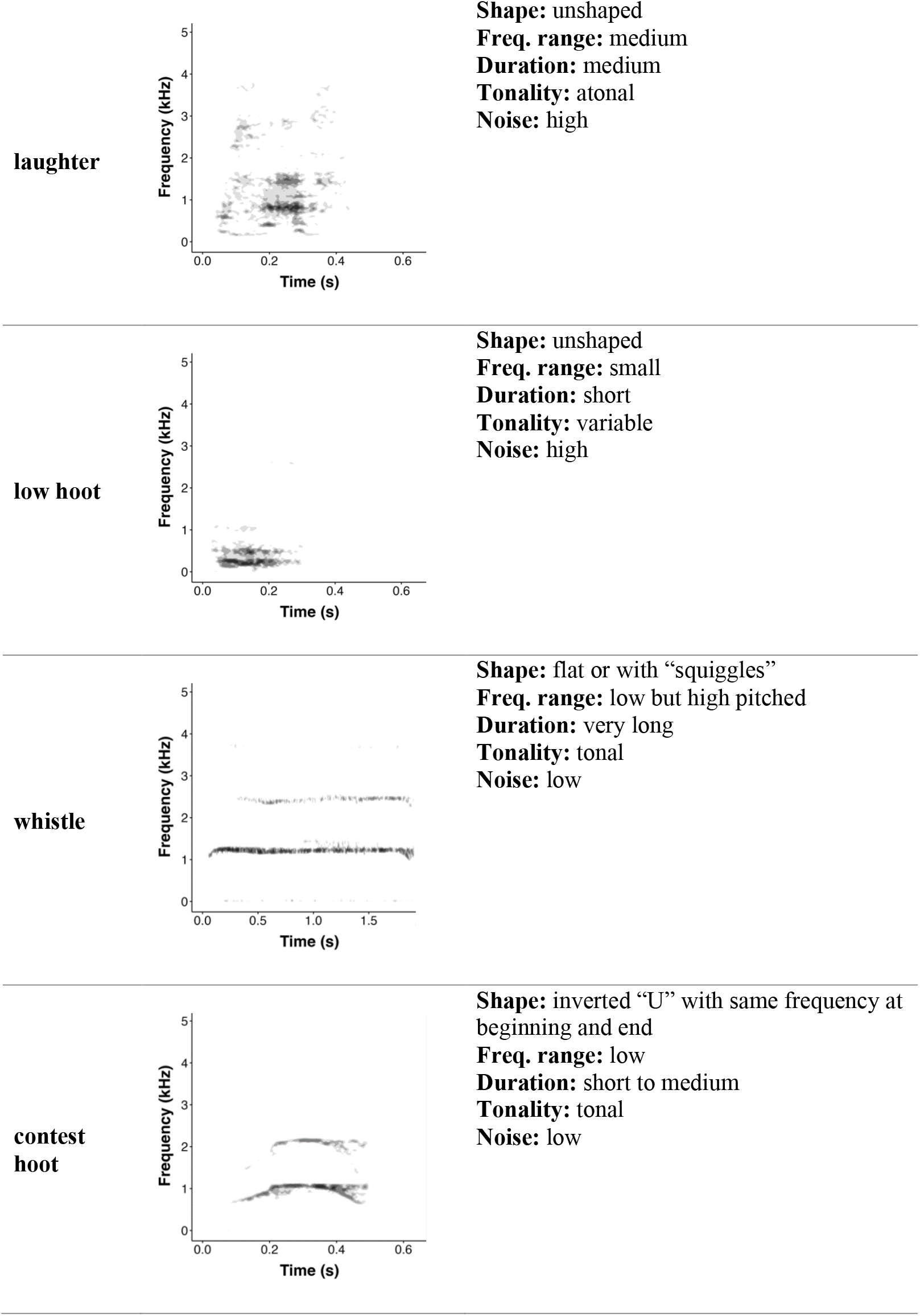
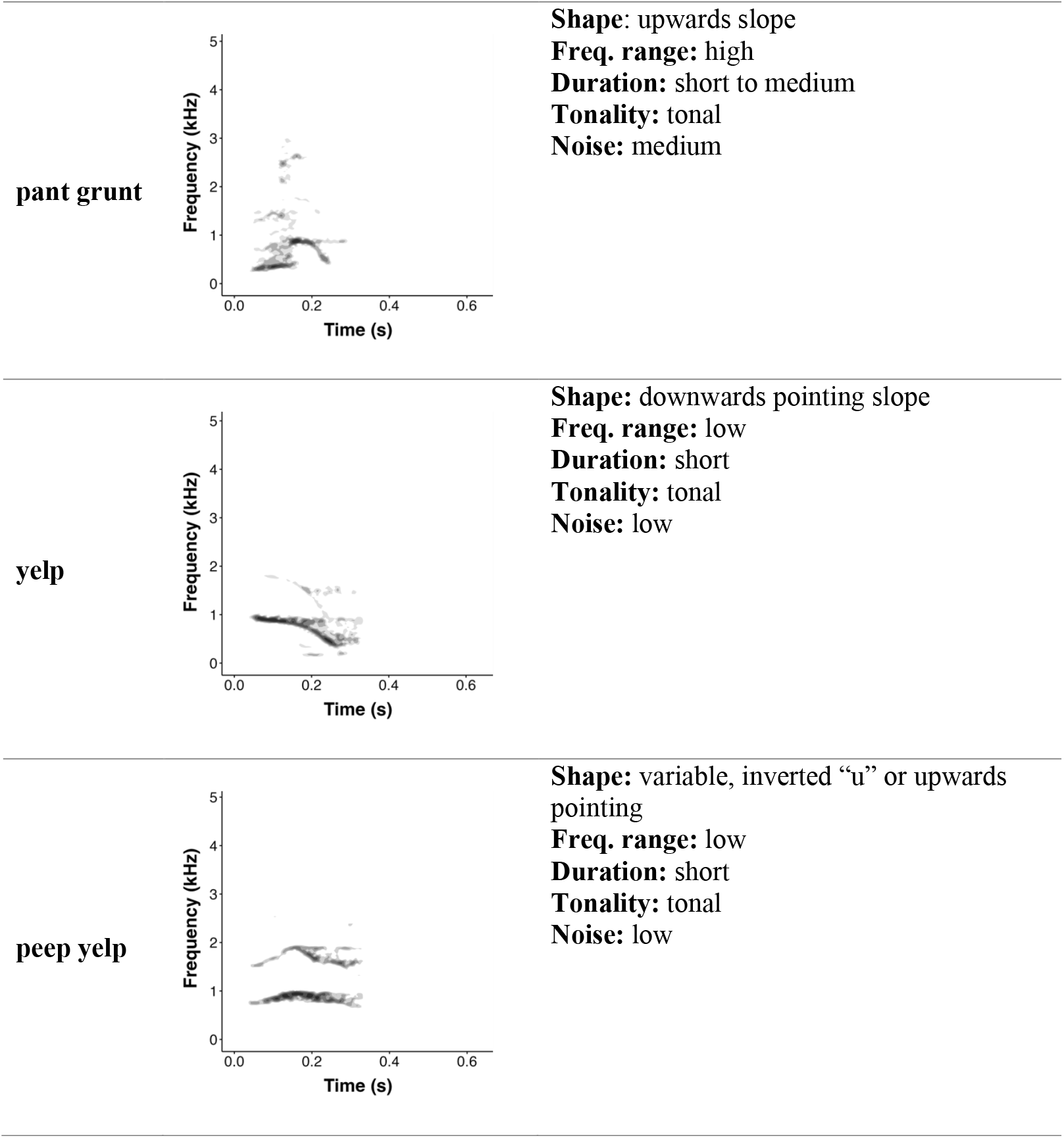
Spectrogram and description of the eleven call types of the updated bonobo vocal repertoire. Note that the x-axes of the spectrograms vary in length. Qualitative identification characteristics are used to describe each call type: “Shape” refers to the form of the fundamental frequency and/or harmonics; “Freq. range” refers to the frequency range of the calls; “Duration” refers to the duration of the call; “Tonality” to the harmonicity of the call; “Noise” to the signal to noise ratio and, if applicable, “Other” refers to additional noteworthy identification characteristics. Spectrograms were made in R with the dynaSpec package (Araya-Salas and Wilkins 2020) with a hanning window, the minimum decibel to be included in the spectrogram set at -30 and silent margins were added at beginning and end. For the spectrograms, corresponding audio recordings can be found in the Electronic Supplementary Material. Spectrograms for the four “bark” variants that are now all classed as high hoots can be found in S5 Fig.

## Discussion

Using supervised random forest analysis, we provide the first quantitative description of the vocal repertoire of wild bonobos. We found that calls can be reliably discriminated into 11 different acoustic categories or vocalisation types ranging from *high hoots* to *grunts, peeps, screams* and *whistles*.

Many of the call types which could be reliably discriminated based on their acoustic structure were also documented by previous research implementing more qualitative approaches. One key difference with existing work, however, is in our proposal to group various *barks* (*scream barks, wieew barks, barks* and *soft barks*) and *high hoots* (with varying degrees of confidence) into a single category. Specifically, we found that all these *barks* were more often classified as *high hoots* than their own call type category (out of bag error rate: *scream barks* (0.73), *wieew barks* (0.74), *barks* (0.76) and *soft barks* (0.95)), suggesting that these four previously discriminated call types rather represent one single call class, here termed *high hoots*. In their qualitative description of the vocal repertoire of bonobos, Bermejo & Omedes (1999) also did not differentiate between *scream barks, wieew barks, high hoots* and *barks*, although they did distinguish between *soft barks* and *barks*.

Where findings also diverge subtly with previous work is with regards to the overall number of comprising call types: de Waal (1980), Bermejo & Omedes (1999) identified 15 and 18 call types, respectively, compared to the 11 clusters we detected. A number of factors might explain these differences. Firstly, our analysis focused exclusively on adult bonobos and hence we excluded additional calls that we recorded from other age classes, such as *pout moans*, which are primarily emitted by immatures. Furthermore, certain rare call types, such as *croaks, hiccups* and *moans* (described by Bermejo & Omedes (1999) could not be included in our dataset due to their highly infrequent occurrence which precluded quantitative analyses, again reducing the number of potential call types that could be acoustically discriminated.

Lastly, whilst the random forest analyses converged on 11 clusters, a considerable degree of acoustic overlap still exists between the call types confirming previous research suggesting the bonobo vocal system is not discrete as is the case in other primate species such as blue monkeys (Fuller 2014), but rather graded, as has also been shown in barbary macaques (*Macaca sylvanus*) (Hammerschmidt and Fischer 1998), lemurs (*Varecia variegata*) (Batist et al. 2023) and indeed in non-primate vocal systems such as dolphins (*Tursiops*) (Jones et al. 2020). Although out of the scope of the current study, follow up research could measure and quantify the extent of gradation for each call type and for the system as a whole within a broader comparative framework. Indeed, recent work has suggested fuzzy clustering, where the degree of gradation of different call types within a repertoire can be assessed (Cusano, Noad, and Dunlop 2021; Wadewitz et al. 2015), to be one key approach that could help to further capture the precise complexity, and with it, the potential flexibility (Keenan et al. 2020) of the bonobo vocal system.

Although quantitative approaches to resolving animal vocal repertoires, such as those implemented here, better avoid the subjective classification of call types, they do not come without their own shortcomings. Specifically, we encountered several obstacles that influenced the quality and analysis of the gathered data that are essential for such “data-hungry” approaches. In line with previous work on similar questions, our dataset can be characterized as SUNG (Small, Unbalanced, Noisy, but Genuine: (Arnaud et al. 2023)) but in addition, we arguably face an even more challenging and SUNG dataset since we compiled data from individuals in their natural habitat, the rainforest, where visibility is inherently lower than in captivity constraining call collection and where unavoidable background noise including cicadas, various birds, and choruses of many other bonobos persists. Background noise in particular represented an important constraint since it made resolving the acoustic parameters for analysis inherently more challenging, ultimately reducing the calls available for follow-up acoustic analyses. A further obstacle encountered in this study was the mandatory 7m distance between humans and bonobos that needed to be adhered to in order to reduce disease transmission and avoid overly-interfering with bonobo behaviour (Gilardi et al. 2015). Whilst this has little influence on higher amplitude long-distance calls such as *high hoots, contest hoots* and *whistles*; softer, short-distance calls we recorded, including *pant grunts* and *grunts* were affected, again representing an additional bottleneck on recordings available for quantification. In addition, our definition for reliably identifying a call type is somewhat arbitrary. Specifically, we used a threshold of 51% - meaning that a call type was considered verified if at least 51% of calls were reliably classified. Future studies could validate the here-presented repertoire particularly when using varying threshold values.

Despite these issues, we are confident our study represents an important first attempt at quantitatively resolving the vocal repertoire of wild bonobos. We hope this work will catalyse similar studies in other primate species where more objective repertoires are still missing including, and particularly surprisingly, chimpanzees. Future work leveraging an even larger sample size could also consider extending our approach to include more unsupervised machine learning-based approaches. Such methods are arguably even more objective since calls are categorized independently of pre-existing call categories further removing observer bias inherent to more supervised approaches.

Lastly, this study provides essential groundwork for follow-up quantitative investigations into the contexts accompanying call types. In particular, a whole repertoire approach can now be adopted to probe how bonobo call types are associated with specific social and environmental events and what light this can shed on their underlying function (see for example Berthet et al., 2023). Such follow-up work will ultimately allow for a more detailed and holistic understanding of bonobo communication.

## Acknowledgements

We thank the pisteurs of the Kokolopori Bonobo Project for their invaluable help with data collection, the Institut Congolais pour la Conservations de la Nature (ICCN) and the Ministry of Scientific Research and Technology in the DRC for their permission to work in the Democratic Republic of the Congo and the Bonobo Conservation Initiative and Vie Sauvage for support. We thank the LuiKotale Bonobo Project for offering access to the study site, the ICCN for granting permission to conduct research in the buffer zone of Salonga National Park, and the people of Lompole village for hosting researchers in their forest. MB thanks Lara Zanutto for help with data collection. IS thanks Claudia Wilke for help with extracting calls. We thank Tim Sainburg for helpful methodological discussions and Nikola Falk for sharing her code to loop the dynaSpec package to create the spectrograms.

## Supporting information

**S1 Table. Sample size of the original call types per individual**

**S2 Table. Acoustic parameters used in the random forest analysis**.

**S3 Analysis. Unsupervised random forest approach**

**S4 Fig. Influence (mean decrease accuracy) each acoustic parameter has on the random forest model**.

**S5 Fig. Spectrograms of four variants of the “high hoot” call type**.

**S6 Table. Acoustic parameter comparison**

