## Supplementary material for "An updated vocal repertoire of wild adult bonobos (*Pan paniscus*)"

**S1 Table. Sample size of the original call types per individual**

| **ID** | **ba** | **bs** | **ch** | **gr** | **hh** | **la** | **lh** | **pe** | **pg** | **py** | **sb** | **sc** | **wb** | **wi** | **ye** | **Total** |
| --- | --- | --- | --- | --- | --- | --- | --- | --- | --- | --- | --- | --- | --- | --- | --- | --- |
| **ID1** | **1** |  |  | **1** |  |  |  | **1** |  |  |  |  |  |  | **2** | **5** |
| **ID2** |  |  |  |  |  |  |  | **2** |  | **3** |  |  |  |  |  | **5** |
| **ID3** |  | **10** | **33** |  | **43** |  | **3** | **5** | **8** | **7** |  |  |  | **2** | **1** | **112** |
| **ID4** |  | **3** |  |  |  |  |  | **2** |  | **2** |  | **24** |  | **3** |  | **34** |
| **ID5** |  |  | **4** |  | **14** |  | **38** |  |  | **6** | **1** |  |  |  |  | **63** |
| **ID6** | **4** | **6** |  |  | **8** |  | **7** | **1** |  | **1** | **2** |  | **1** |  |  | **30** |
| **ID7** |  |  | **9** |  |  |  |  |  |  |  | **1** |  |  |  |  | **10** |
| **ID8** | **2** |  |  |  |  |  |  |  |  | **2** |  |  |  | **5** |  | **9** |
| **ID9** |  |  |  | **2** |  |  |  | **1** |  |  |  |  |  |  | **2** | **5** |
| **ID10** |  |  |  |  |  |  |  |  |  |  | **1** |  |  |  |  | **1** |
| **ID11** |  | **1** |  | **2** | **3** |  |  | **1** |  | **1** |  | **5** | **2** | **1** |  | **16** |
| **ID12** | **3** |  |  | **1** | **4** |  |  |  |  |  | **1** |  |  | **1** |  | **10** |
| **ID13** |  |  | **3** |  |  |  |  |  |  |  |  |  |  | **1** |  | **4** |
| **ID14** | **6** | **1** |  |  | **5** |  |  | **3** |  | **1** | **6** | **1** |  |  |  | **23** |
| **ID15** |  |  |  | **1** |  |  |  | **1** |  | **1** |  |  |  |  |  | **3** |
| **ID16** |  |  |  |  |  |  |  | **2** |  |  |  |  |  | **1** |  | **3** |
| **ID17** |  |  |  | **1** |  |  |  |  |  | **1** |  |  |  |  |  | **2** |
| **ID18** |  |  |  |  | **1** |  |  | **6** |  | **4** | **4** | **1** |  |  | **17** | **33** |
| **ID19** | **16** | **1** | **2** | **15** | **49** |  | **7** | **26** |  | **7** | **10** | **1** | **2** | **1** | **8** | **145** |
| **ID20** |  |  |  |  |  |  |  |  |  | **1** |  |  |  |  |  | **1** |
| **ID21** |  |  |  | **2** | **41** |  | **3** | **7** |  | **11** | **10** |  | **31** | **1** | **10** | **116** |
| **ID22** |  |  |  | **6** | **7** |  |  | **1** |  |  |  |  | **2** |  | **2** | **18** |
| **ID23** | **13** | **3** |  | **24** |  |  | **10** | **4** |  | **12** | **9** | **8** | **5** | **3** | **2** | **93** |
| **ID24** |  |  | **2** |  |  |  |  |  |  | **6** |  |  |  |  |  | **8** |
| **ID25** |  |  |  |  |  |  |  |  |  |  | **1** |  |  |  |  | **1** |
| **ID26** |  |  |  |  |  |  | **1** | **1** |  | **1** |  |  |  | **4** |  | **7** |
| **ID27** |  |  |  | **4** |  |  |  | **4** |  |  |  |  |  |  |  | **8** |
| **ID28** |  |  |  |  | **8** |  |  |  |  |  |  |  |  |  |  | **8** |
| **ID29** |  |  |  |  | **6** | **6** |  |  |  |  |  | **9** |  |  |  | **21** |
| **ID30** | **8** | **12** |  | **2** | **10** |  |  | **8** |  | **5** | **7** | **4** |  | **19** |  | **75** |
| **ID31** | **6** |  |  | **9** | **1** |  | **2** | **2** |  | **8** |  | **1** |  | **1** | **1** | **31** |
| **ID32** |  |  |  | **1** | **3** |  |  | **4** | **1** | **1** | **1** |  |  |  |  | **11** |
| **ID33** |  |  |  | **17** |  |  |  | **17** |  | **4** |  |  |  | **5** | **10** | **53** |
| **ID34** |  |  |  |  | **1** |  |  |  | **1** |  |  |  | **1** |  |  | **3** |
| **ID35** | **6** |  |  |  |  |  |  |  |  | **2** | **2** | **1** |  |  |  | **11** |
| **ID36** | **4** |  |  |  |  |  |  |  |  | **2** | **1** |  |  |  |  | **7** |
| **ID37** |  |  |  | **1** |  |  |  | **4** |  | **8** | **3** |  |  |  | **2** | **18** |
| **ID38** |  |  |  |  |  |  |  |  |  |  |  |  |  | **7** |  | **7** |
| **ID39** | **1** |  |  | **1** |  |  |  |  |  | **2** | **1** |  |  |  | **2** | **7** |
| **ID40** |  |  |  | **1** |  |  |  | **2** |  |  | **2** | **2** |  |  | **2** | **9** |
| **ID41** | **1** |  |  | **2** |  |  |  | **2** |  | **2** |  |  | **1** | **1** | **2** | **11** |
| **ID42** |  |  |  | **5** |  |  |  |  |  |  |  |  |  |  |  | **5** |
| **ID43** | **4** |  |  |  | **2** |  |  |  |  |  |  |  | **1** | **1** |  | **8** |
| **ID44** |  |  |  |  | **1** |  |  |  |  |  |  |  |  |  |  | **1** |
| **ID45** |  |  |  |  | **6** |  |  |  |  |  |  |  |  | **1** |  | **7** |
| **ID46** | **6** | **1** |  |  | **21** |  |  | **2** |  |  | **1** | **12** |  | **5** | **1** | **49** |
| **ID47** |  |  |  | **29** | **7** |  |  | **7** | **7** | **31** | **3** | **1** | **1** | **7** | **12** | **105** |
| **ID48** | **6** |  |  | **1** | **4** |  |  | **3** |  | **1** |  |  |  | **6** | **6** | **27** |
| **ID49** |  |  | **8** |  |  |  |  |  |  |  |  |  |  |  |  | **8** |
| **ID50** | **16** |  |  |  | **23** |  |  | **10** | **3** | **11** | **3** | **7** | **9** | **9** | **8** | **99** |
| **ID51** | **1** |  |  | **7** | **1** |  |  | **12** |  | **1** | **7** |  |  | **1** | **8** | **38** |
| **ID52** |  | **2** |  |  | **7** | **23** | **1** | **3** | **18** | **7** |  | **5** |  | **1** |  | **67** |
| **ID53** |  |  |  |  | **9** |  | **4** | **7** | **15** | **2** | **2** | **4** | **2** | **4** | **9** | **58** |
| **Total** | **104** | **40** | **61** | **135** | **285** | **29** | **76** | **151** | **53** | **154** | **79** | **86** | **58** | **91** | **107** | **1509** |

**S2 Table. Acoustic parameters used in the random forest analysis.**

| ***Abbreviation*** | ***Name*** | ***Description*** |
| --- | --- | --- |
| duration | call duration | length of the call (s) |
| ***meanfreq*** | mean frequency | Mean of frequency spectrum (i.e. weighted average of frequency by amplitude within supplied band pass) (in kHz) |
| **sd** | standard deviation | standard deviation of frequency weighted by amplitude |
| **freq.median** | median frequency, kHz | The frequency at which the signal is divided in two fre- quency intervals of equal energy (in kHz) |
| **freq.Q25** | first quantile of frequency | The frequency at which the signal is divided in two fre- quency intervals of 25% and 75% energy respectively (in kHz) |
| **freq.Q75** | third quantile of frequency | The frequency at which the signal is divided in two frequency intervals of 75% and 25% energy respectively (in kHz) |
| **freq.IQR** | interquartile frequency range | Frequency range between ’freq.Q25’ and ’freq.Q75’ (in kHz) |
| **time.median** | median time | The time at which the signal is divided in two time intervals of equal energy (in s) |
| **time.Q25** | first quartile time | The time at which the signal is divided in two time intervals of 25% and 75% energy respectively (in s). |
| **time.Q75** | third quartile time | The time at which the signal is divided in two time intervals of 75% and 25% energy respectively (in s). |
| **time.IQR** | interquartile time range | Time range between ’time.Q25’ and ’time.Q75’ (in s). |
| **skew** | skewness | Asymmetry of the spectrum |
| **kurt** | kurtosis | Peakedness of the spectrum |
| **sp.ent** | spectral entropy | Energy distribution of the frequency spectrum. Pure tone ~ 0; noisy ~ 1. |
| **time.ent** | time entropy | Energy distribution on the time envelope. Pure tone ~ 0; noisy ~ 1. |
| **entropy** | entropy | Product of time and spectral entropy sp.ent * time.ent. |
| **sfm** | spectral flatness | Similar to sp.ent (Pure tone ~ 0; noisy ~ 1). |
| **meandom** | average dominant frequency |  |
| **mindom** | minimum dominant frequency |  |
| **maxdom** | maximum dominant frequency |  |
| **dfrange** | range of dominant frequency measured across the acoustic signal |  |
| **modindx** | modulation index | Calculated as the cumulative absolute difference between ad- jacent measurements of dominant frequencies divided by the dominant frequency range. 1 means the signals is not modulated. |
| **startdom** | dominant frequency measurement at the start of the signal |  |
| **enddom** | dominant frequency measurement at the end of the signal |  |
| **dfslope** | slopeofthechangeindominantfrequencythroughtime((enddom-startdom)/duration). Units are kHz/s. |  |
| **meanpeakf** | average peak frequency | Frequency with highest energy from the mean frequency spectrum. Typically, more consistent than peakf. |
| **dtw.dim.1** | dynamic time warping |  |
| **dtw.dim.2** | dynamic time warping |  |
| **dtw.dim.3** | dynamic time warping |  |
| **dtw.dim.4** | dynamic time warping |  |
| **dtw.dim.5** | dynamic time warping |  |

*The abbreviation of the parameter (which is used in S1 Fig), the full name and a description of the parameter are given. This table is partly taken from (Keen et al. 2021).*

**S3 Analysis. Unsupervised random forest approach**

An unsupervised random forest approach yielded in only two clusters / different call types.

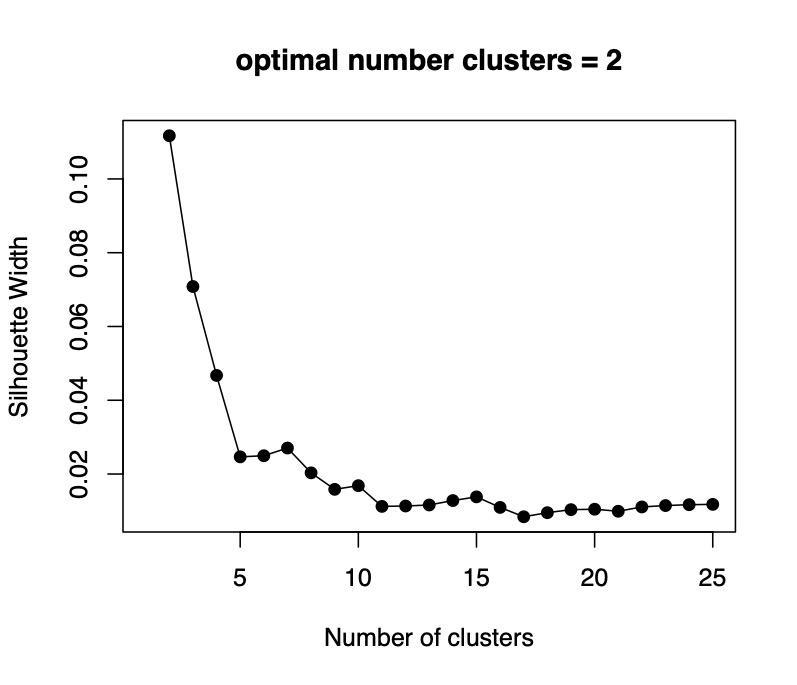

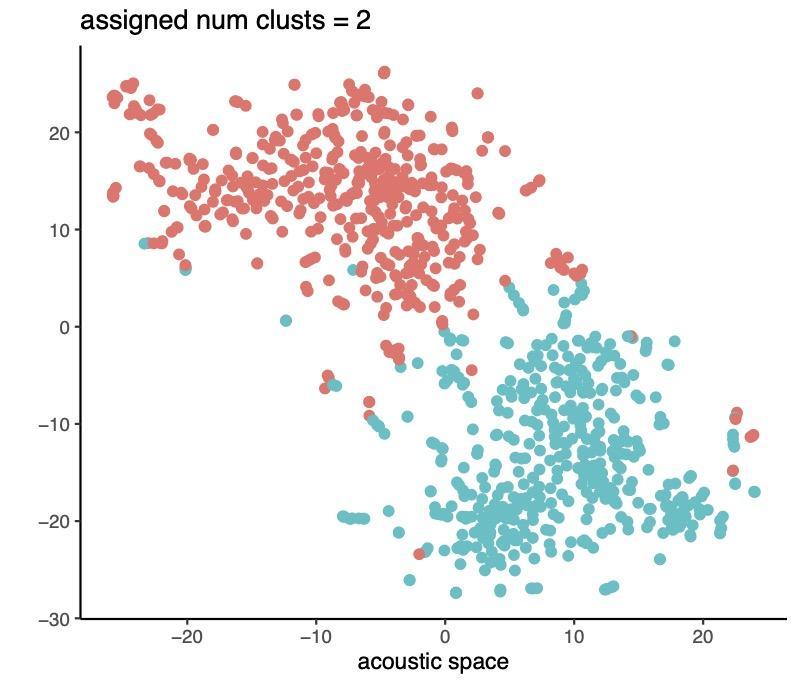

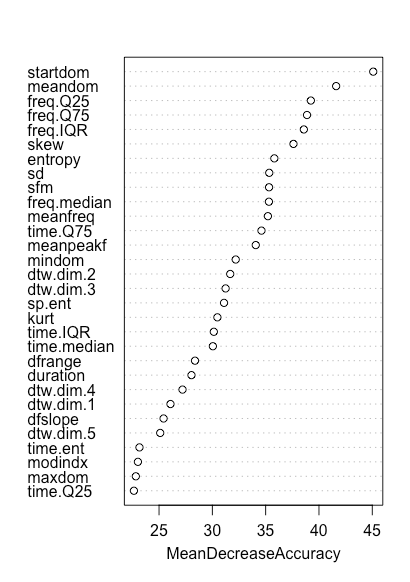

**S4 Fig. Influence (mean decrease accuracy) each acoustic parameter has on the random forest model.** See S2 Table for full name and description of the acoustic parameters.

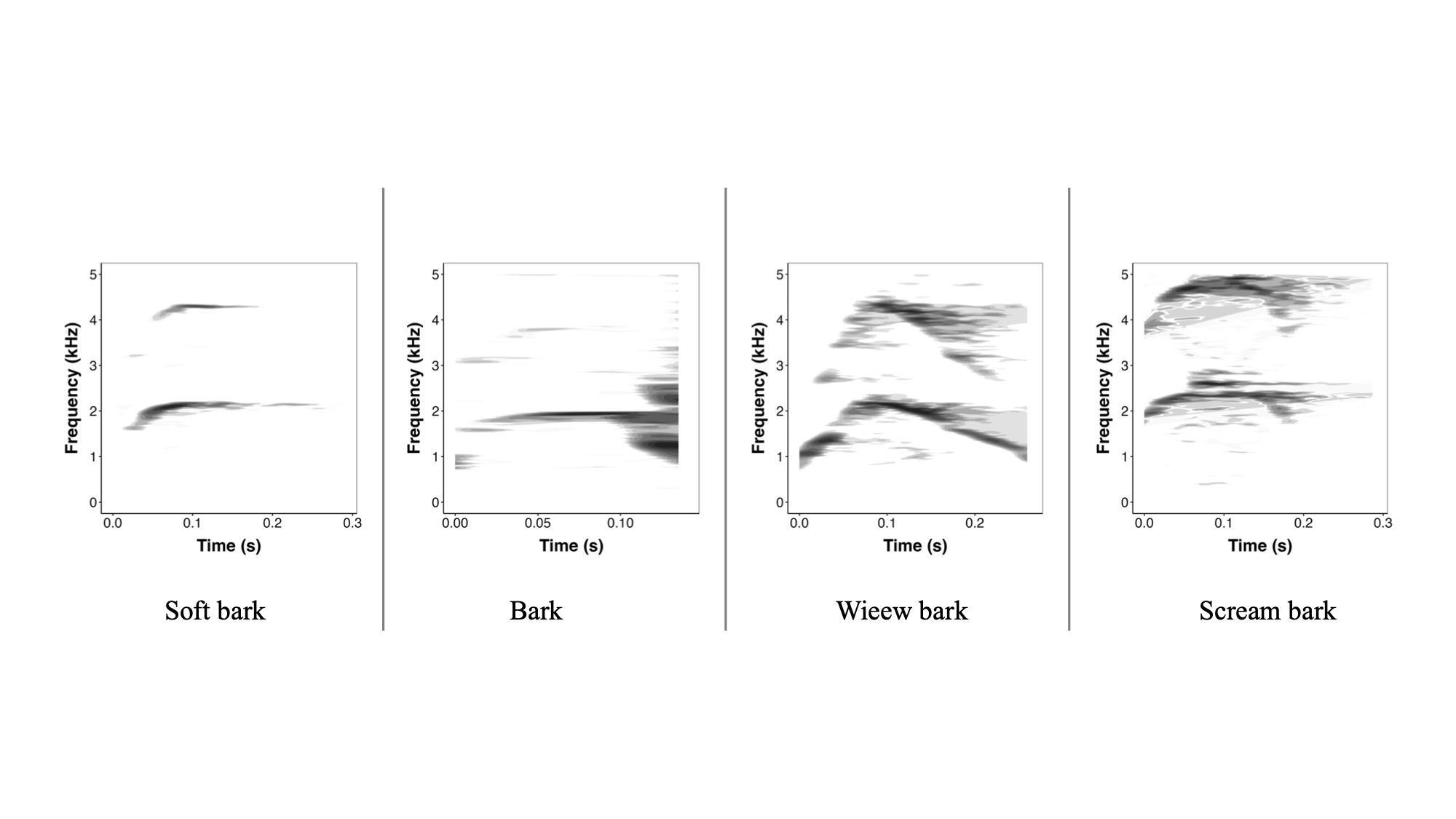

**S5 Fig. Spectrograms of four variants of the “high hoot” call type.**  These calls were formerly, as “original call types”, categorized as “soft bark”, “bark”, “wieew bark” and “scream bark”. Using our random forest analysis, these former call types are now merged together into a single call category: “high hoots”.

**S6 Table. Acoustic parameter comparison**

| **Parameter** | **Paramater Description** | **Wegdell et al.** | **Keenan et al.** | **Arnaud et al.** |
| --- | --- | --- | --- | --- |
| call duration | length of the call (s) | X | X | X |
| mean frequency | Mean of frequency spectrum (i.e. weighted average of frequency by amplitude within supplied band pass) (in kHz) | X |  |  |
| standard deviation | standard deviation of frequency weighted by amplitude | X |  |  |
| median frequency, kHz | The frequency at which the signal is divided in two fre- quency intervals of equal energy (in kHz) | X | X | X |
| first quantile of frequency | The frequency at which the signal is divided in two fre- quency intervals of 25% and 75% energy respectively (in kHz) | X | X | X |
| third quantile of frequency | The frequency at which the signal is divided in two frequency intervals of 75% and 25% energy respectively (in kHz) | X | X | X |
| interquartile frequency range | Frequency range between ’freq.Q25’ and ’freq.Q75’ (in kHz) | X |  |  |
| median time | The time at which the signal is divided in two time intervals of equal energy (in s) | X | X | X |
| first quartile time | The time at which the signal is divided in two time intervals of 25% and 75% energy respectively (in s). | X |  | X |
| third quartile time | The time at which the signal is divided in two time intervals of y75% and 25% energy respectively (in s). | X |  | X |
| interquartile time range | Time range between ’time.Q25’ and ’time.Q75’ (in s). | X |  |  |
| skewness | Asymmetry of the spectrum | X |  |  |
| kurtosis | Peakedness of the spectrum | X |  |  |
| spectral entropy | Energy distribution of the frequency spectrum. Pure tone ~ 0; noisy ~ 1. | X |  |  |
| time entropy | Energy distribution on the time envelope. Pure tone ~ 0; noisy ~ 1. | X |  |  |
| entropy | Product of time and spectral entropy sp.ent * time.ent. | X |  |  |
| spectral flatness | Similar to sp.ent (Pure tone ~ 0; noisy ~ 1). | X |  |  |
| average dominant frequency |  | X |  |  |
| minimum dominant frequency |  | X |  |  |
| maximum dominant frequency |  | X | X |  |
| range of dominant frequency measured across the acoustic signal |  | X |  |  |
| modulation index | Calculated as the cumulative absolute difference between ad- jacent measurements of dominant frequencies divided by the dominant frequency range. 1 means the signals is not modulated. | X |  |  |
| dominant frequency measurement at the start of the signal |  | X |  |  |
| dominant frequency measurement at the end of the signal |  | X |  |  |
| slopeofthechangeindominantfrequencythroughtime((enddom-startdom)/duration). Units are kHz/s. |  | X |  |  |
| average peak frequency | Frequency with highest energy from the mean frequency spectrum. Typically, more consistent than peakf. | X |  |  |
| dynamic time warping |  | X |  |  |
| dynamic time warping |  | X |  |  |
| dynamic time warping |  | X |  |  |
| dynamic time warping |  | X |  |  |
| dynamic time warping |  | X |  |  |
| Harmonics-to-noise ratio (dB) |  |  |  | X |
| Maximum fundamental frequency reached over the call (Hz) |  |  | X | X |
| relative time at which f.max is reached (%) |  |  |  | X |
| Fundamental frequency at the beginning of the call (Hz) |  |  | X | X |
| Fundamental frequency at the temporal middle of the call (Hz) |  |  |  | X |
| Fundamental frequency at the end of the call (Hz) |  |  | X | X |
| Average fundamental frequency over the call (Hz) |  |  |  | X |
| Raw fundamental frequency slope (start-middle), in Hertz |  |  |  | X |
| Raw fundamental frequency slope (middle-end), in Hertz |  |  |  | X |
| dct0 DCT coefficient |  |  |  | X |
| dct1 DCT coefficient |  |  |  | X |
| dct2 DCT coefficient |  |  |  | X |
| dct3 DCT coefficient |  |  |  | X |
| dct4 DCT coefficient |  |  |  | X |
| Average value of the Mel-frequency cepstral coefficients (MFCCs) over the call (unitless) Average value of the 𝛥MFCCs over the call (unitless) |  |  |  | X |
| Average value of the 𝛥𝛥MFCCs over the call (unitless) |  |  |  | X |
| Deviation of the MFCCs over the call (unitless) |  |  |  | X |
| Average value of the Mel-frequency cepstral coefficients (MFCCs) over the call (unitless) Average value of the 𝛥MFCCs over the call (unitless) |  |  |  | X |
| Deviation of the 𝛥MFCCs over the call (unitless) MFCC |  |  |  | X |
| Deviation of the 𝛥𝛥MFCCs over the call (unitless) |  |  |  | X |
| F0-Peak Time | Point over the duration of the call at which F0-Peak is reached. Manually calculated as a proportion: time of F0-Peak(s)/Call Duration (s) |  | X |  |
| Ascending Slope | Calculated as: (F0-Peak e F0-Start)/(F0-Peak Time e 0) |  | X | X |
| Descending Slope | Calculated as: (F0-End e F0-Peak)/(1 e F0-Peak Time) |  | X | X |
| Slope: F0-Start to Midpoint | Calculated as: (F0 at midpoint of call duration e F0-Start)/(Time at midpoint of call duration e 0) |  | X | X |
| Slope: F0-Midpoint to End | Calculated as: (F0-End e F0 at midpoint of call duration)/(Call duration e Time at midpoint of call duration) |  | X | X |
| Maximum Time | The ﬁrst time point along the call where maximum amplitude occurs on waveform (s) |  | X |  |

The two hitherto performed quantitative analyses of a subset of calls of the vocal repertoire of bonobos (Arnaud et al. 2023 and Keenan et al. 2020) and our study used broadly similar, commonly used acoustic parameters for the quantitative acoustic analyses. All three studies used parameters such as duration and parameters related to the distribution of the energy within the call. Whilst we in the study at hand analysed acoustic parameters related to the dominant frequency, Arnaud et al. and Keenan et al. used acoustic features regarding the fundamental frequency. Oftentimes, but not always, the dominant frequency correlates highly with the fundamental frequency. In addition, whilst we used dynamic time warping-related parameters, Arnaud et al. used MFCCs, and Keenan used neither of the two.
